# Motor Sequences - Separating The Sequence From The Motor. A longitudinal rsfMRI Study

**DOI:** 10.1101/2021.02.09.430495

**Authors:** ATP Jäger, JM Huntenburg, SA Tremblay, U Schneider, S Grahl, J Huck, CL Tardif, A Villringer, CJ Gauthier, PL Bazin, CJ Steele

## Abstract

In motor learning, sequence-specificity, i.e. the learning of specific sequential associations, has predominantly been studied using task-based fMRI paradigms. However, offline changes in resting state functional connectivity after sequence-specific motor learning are less well understood. Previous research has established that plastic changes following motor learning can be divided into stages including fast learning, slow learning and retention. A description of how resting state functional connectivity after sequence-specific motor sequence learning (MSL) develops across these stages is missing. This study aimed to identify plastic alterations in whole-brain functional connectivity after learning a complex motor sequence by contrasting an active group who learned a complex sequence with a control group who performed a control task matched for motor execution. Resting state fMRI and behavioural performance were collected in both groups over the course of 5 consecutive training days and at follow-up after 12 days to encompass fast learning, slow learning, overall learning and retention. Between-group interaction analyses showed sequence-specific decreases in functional connectivity during overall learning in the right supplementary motor area (SMA). We found that connectivity changes in key regions of the motor network including the superior parietal cortex (SPC) and primary motor cortex (M1) were not a result of sequence-specific learning but were instead linked to motor execution. Our study confirms the sequence-specific role of SMAthat has previously been identified in online task-based learning studies, and extends it to resting state network changes after sequence-specific MSL.

## Introduction

Motor learning induced brain plasticity has been typically studied using magnetic resonance imaging (MRI) (for review, see Dayan & Cohen, 2011; Krakauer et al, 2019; Taubert et al., 2012), from immediate online functional changes (Coynel et al., 2010, Karim et al., 2017, Steele & Penhune, 2010; Yokoi & Diedrichsen, 2019) to long term structural effects (Gryga et al., 2012, Taubert et al., 2010, Bengtsson et al., 2005; Scholz et al., 2009). While task-based studies investigate immediate training-related processes during task performance and structural studies usually identify longer term changes in brain morphometry they cannot comment on how the learned skill is represented and maintained within functional networks of the brain outside of the training/learning context. Resting state fMRI (rsfMRI) can be used to investigate functional plasticity that occurs between online functional and slower structural changes. Measured in the absence of a task, resting-state network dynamics are thought to reflect the previous co-activation of functionally connected brain regions (Biswal et al., 1994, Guerra-Carrillo et al., 2014). Alterations in these functional networks are thought to reflect the strengthening of the memory trace generated during practice (Albert et al, 2009, Lewis et al., 2009, Vahdat et al., 2011). Therefore, assessing resting state networks and how they change in response to training can provide unique insight into training-related functional plasticity beyond the immediate time point of learning.

Previous research in motor learning has established that sequence learning *per se* can be distinguished from motor execution (Rosenbaum et al., 1983, Penhune and Steele, 2012, Wiestler & Diedrichsen, 2013). However, while this concept has been assessed, tested and discussed in task-based studies (Seidler et al., 2002; Yokoi and Diedrichsen, 2019, Wymbs & Grafton, 2015), learning that is specific to sequential associations (i.e., sequence-specific learning) is rarely if ever differentiated from motor execution in rsfMRI studies of motor learning. In part, this is due to the added challenge of including additional control groups matched for motor execution. Conclusions about functional changes being the result of sequence-specific learning, and not simply an effect of repeated performance, cannot be made without the inclusion of a control group (Thomas and Baker, 2013, Steel et al. 2019). As a result, it is currently unclear which regions of the sensorimotor network are involved in offline sequence-specific learning versus those more generally involved in motor execution.

RsfMRI connectivity changes after motor sequence learning (MSL) have been investigated with “pre/post” frameworks where resting state functional connectivity (rsFC) is typically measured only before and/or after performing the task to identify immediate and overnight effects (Sami and Miall, 2013, Gregory et al., 2014, Sami et al., 2014, Mary et al., 2017, Albert et al., 2009). And while motor learning is thought to progress through several stages involving rapid changes in performance (fast learning), slower improvements over a longer time period (slow learning), and maintenance of robust performance (retention) (Doyon & Benali, 2005, Dayan and Cohen, 2011, Lohse et al., 2014), there are few studies that have followed changes in rsfMRI networks after MSL across multiple sessions of training to investigate larger timescales (Ma et al., 2011, Xiong et al., 2009). To our knowledge, there has been no investigation of sequence-specific resting state network plasticity across all three stages of learning. The present study is the first to address this gap.

Here, participants learned a continuous motor task with their right (dominant) hand over five days of training, followed by a retention probe 12 days later. RsfMRI was collected five times over the course of the training, providing a rich sampling of the different stages of learning in the absence of performance. Crucially, participants were randomly assigned to either a training (sequence-specific training) or control group (matched for motor execution). This rich dataset allowed us to identify the brain regions involved in sequence-specific learning and explore their functional dynamics.

## Methods

### Participants

40 right-handed healthy participants with no history of neurological disorder (22 females, ranging from 20 to 32 years of age, M ± SD: 24.5 ± 2.44) were included in this study. Participants were recruited from the database of the Max Planck Institute for Human Cognitive and Brain Sciences in Leipzig, Germany and randomly assigned to an experimental (N=20, 11 females) and control group (N=20, 10 females). The experimental group learned a previously published complex visuomotor sequence pinch-force task (Gryga et al., 2012) while the control group performed a much simpler sinusoidal sequence that required almost no learning to perform. The training period consisted of 5 days, with an additional familiarization session prior to training and a retention session following 12 days without practice. Participants did not meet any MRI exclusion criteria, gave written informed consent in accordance with the Declaration of Helsinki, and were monetarily compensated for their time. The study design was approved by the ethics review board of the Medical Faculty of the University of Leipzig.

### Task and Stimuli: Sequential Pinch Force Task

The Sequential Pinch Force Task (SPFT) is a motor sequence task that requires the participant to use the index finger and thumb of the dominant (right) hand to exert force on the end of a pinch force device (Fig. 1a) which measures force at a sample rate of 80Hz. This pinching force controls the height of a rectangular yellow force bar (FOR) that is displayed on the screen (Fig. 1b).

**Fig. 1.**
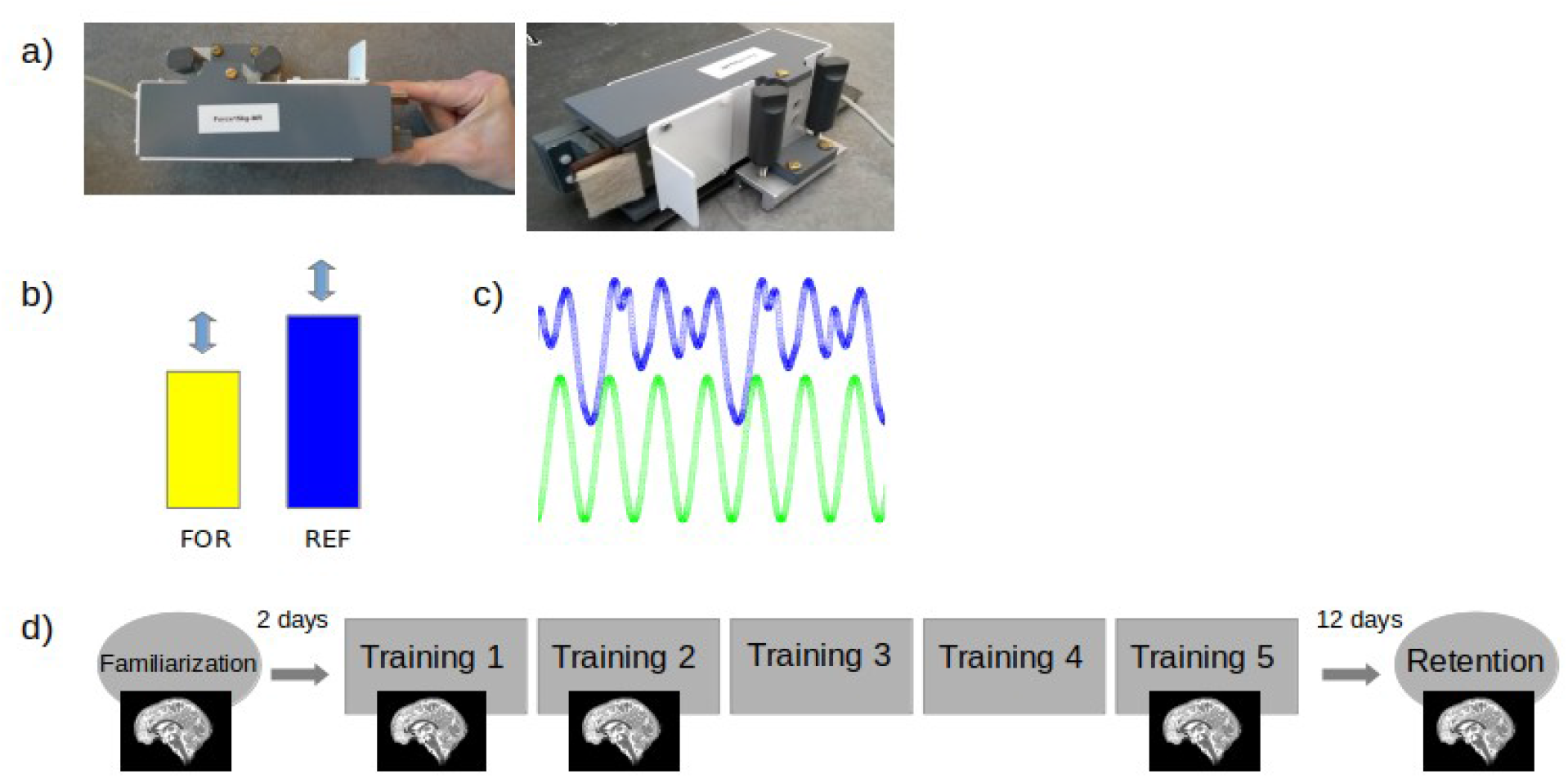
Task and Procedure. a) Sequential Pinch Force Task (SPFT) device, b) Visual screen representations: FOR bar (yellow) REF bar (blue), c) Pinch-force task, green: simple control condition (SMP) sequence, blue: learning condition (LRN) sequence d) experimental design. Brain images indicate sessions where 7T scanning was performed.

In the SPFT, the participant is instructed to adjust the height of FOR to match that of an adjacent blue reference (REF) bar. The up and down movement of REF follows a pre-set sequence (Figure 1b, 1c). There are 3 conditions: a learning condition (LRN), a simple control condition (SMP) and a resting condition (RST). In LRN, the movement of the bar follows a previously published sequence of varying heights that is difficult to predict (Fig. 1c) and learned over time (Gryga, 2012), while the SMP sequence is a simple sinusoid. The SMP sequence was designed to match the LRN sequence for its frequency of maximum power, duration, range, and the total magnitude of force. While in the SMP condition participants merely performed this predictable and unvarying sinusoidal sequence of motor movements; in the LRN condition participants execute the same *type* of movements (i.e., varying pinch force over time) but performed a more complex sequence (Figure 1c). As a result, SMP serves as a matched control for motor execution which, in comparison with LRN, can be used to identify sequence-specific differences that are over and above any changes associated with motor execution. During RST, both bars are displayed statically at 50% of their maximum height and participants were instructed to focus their gaze between the upper edges of the two bars.

On each training day, 3 blocks consisting of 3 SMP, LRN and RST trials each were performed by the experimental group; for a total of 9 trials per condition on each day. In order to ensure that the training was the same on all days, the order was fixed across participants such that they started with the SMP trials followed by RST and then LRN trials. The control group performed the same number of blocks and trials but never performed LRN: the LRN blocks were replaced by SMP blocks such that the total effort for each group remained the same. Each trial was 18 seconds in length. The sequential force trace was the same across participants to ensure that all participants faced the same sequence difficulty.

After the SMP and LRN trials, participants received feedback reflecting their average temporal accuracy in matching the heights of REF and FOR. The average temporal accuracy consisted of the mean time lag per block (in ms).

For simplicity, we will refer to the groups based on which sequence they were assigned throughout the rest of this paper: SMP (SMP only) vs. LRN (SMP and LRN).

### Training and Experimental Procedure

The experimental procedure (Fig. 1d) consisted of one familiarization session (d0), five training sessions (d1-d5) and one retention session (d17, which took place 12 days after d5). The first training session (d1) always took place on Monday and the d0 familiarization session took place on the previous Friday so that the training period for each participant always took place on five consecutive days (Monday-Friday). The familiarization session included a maximum force calibration step followed by 9 trials of the SMP sequence. When measuring the maximum pinch force, the subject was asked to pinch the device with as much force as possible for ten seconds and the maximum value obtained during this period was considered maximum force. This measurement was used to calibrate the level of force required to move the visually presented force bar and was set to between 5% (minimum bar level) and 30% (maximum bar level) of each individual’s maximum force to ensure that participants were also matched by relative effort. Five sessions (d0, d1, d2, d5, d17) were performed while lying supine inside of the MRI scanner and two (d3, d4) outside of the scanner in a separate testing room while seated at a computer. During these sessions, the SPFT was performed with the dominant hand, over a total duration of 20 minutes (3 blocks, 9 trials [LRN group: 3x(LRN), 3x(SMP), 3x(RST); SMP group: 3x(SMP), 3x(SMP), 3x (RST)]). The d17 retention session followed the same protocol as the training days and was measured 12 days after the last day of the training period. All sessions took place at the same time during the day to account for potential time of day fluctuations in resting state connectivity (Steel et al. 2019).

### MRI protocol

MRI data were acquired on a 7 Tesla MRI scanner (MAGNETOM, Siemens Healthcare, Erlangen, Germany) equipped with a 32-channel head coil (Nova). For the purposes of the current study, blood-oxygenation-level dependent (BOLD) rsfMRI, MP2RAGE T1 (Marques et al., 2010), and a fieldmap were acquired. RsfMRI scans [BOLD, voxel dimensions = 1.2 x 1.2 x 1.2mm, 512 whole brain volumes, FOV = 192 x 192 mm², slice acceleration factor: 2, slice thickness = 1mm, 102 slices, GRAPPA factor 2, partial Fourier 6/8, TR = 1130ms, TE = 22ms, flip angle = 40°, bandwidth = 1562 Hz/Px] were acquired under an eyes open condition with a fixation cross for 10 minutes and took place before the task. Additionally, MP2RAGE images [voxel dimensions = 0.7 x 0.7 x 0.7mm, FOV = 224 x 224 x 240mm³, TR = 5000ms, TE = 2.45ms, flip angle 1 = 5°, flip angle 2 = 3°, bandwidth = 250 Hz/Px] and a fieldmap [voxel dimensions = 2 x 2 x 2mm, FOV = 256 x 256mm², slice thickness = 2mm, 80 slices, TR = 18ms, TE1 = 4.08ms, TE2 = 9.18ms, flip angle = 10°, bandwidth = 300 Hz/Px] were also collected.

### Image Processing

A custom preprocessing pipeline was implemented using Nipype (Gorgolewski et. al. 2011) and Nilearn (v0.2.3, Abraham et al., 2014). The first five volumes of the rsfMRI EPI sequences were removed for signal stabilization. Motion correction was performed using the SpaceTimeRealign algorithm (Roche, 2011) implemented in Nipy. Given the short TR, no slice timing correction was applied. Magnetic field inhomogeneity and subsequent image distortions are more pronounced at higher field strengths (Cusack et al., 2003), therefore EPI data were undistorted using the fieldmaps and FSL FUGUE (Jenkinson, 2004). Outliers in composite motion and average brain signal were detected using Nipype’s ArtifactDetect function for removal in a subsequent step. Nilearn’s high_variance_confounds function was used to extract time courses representative of physiological noise from the white matter and cerebrospinal fluid, following the aCompCor approach (Behzadi et al., 2007). Nuisance regression was performed using Nilearn’s NiftiMasker and included the previously calculated 12 motion regressors (3 translations and 3 rotations plus their first derivatives), outlier regressors and physiological regressors. Timeseries were detrended, variance-normalized and bandpass filtered between 0.01-0.1 Hz. A 2.4 mm FWHM Gaussian spatial smoothing filter was applied to the preprocessed data (Poldrack et al., 2011). Eigenvector centrality (EC) maps were calculated with the fast Eigenvector centrality mapping (fastECM) algorithm by Wink et al., (2012) in native space to protect data properties. Finally, a linear (12 degree of freedom (dof)) and nonlinear (Syn) registration to MNI space were performed with ANTS (Avants et al., 2009) as implemented in the CBS Tools (Bazin et al., 2014). Further preprocessing steps such as generating transformation maps between native, group and MNI spaces, as well as tissue segmentation of the anatomical MP2RAGE images were performed using the CBS Tools. All scripts are openly available at https://github.com/AthSchmidt/MMPI/tree/master/preprocessing.

### Network Analysis

In recent years, graph-based analyses have become increasingly used for studying functional connectivity (Sami and Miall, 2013, Wang, 2010 review). In such analyses, brain regions are treated as nodes – and several approaches have been used to describe the dynamics of connections between such nodes (Bullmore and Bassett, 2011). Graph based analyses also offer the possibility to undertake data-driven investigations of brain dynamics globally and quantifiably on a voxel level without prior assumptions. . EC mapping is a method to analyze network structures which identifies nodes that are of central importance to the entire network (Lohmann et al., 2010). The importance of nodes is determined by the connectedness to other nodes, increasing as the connection to nodes with many important connections also increases. In other words: nodes become more important to the network when they are connected to other important nodes, and are assigned higher EC values. The weighting of the nodes is achieved by calculating their eigenvector from a similarity matrix. (Lohmann et al., 2010). The dominant eigenvector is the one with the largest eigenvalue of this matrix. The coefficient in this dominant eigenvector is computed iteratively as the weighted sum of centralities of the neighbors of a given node (Wink, 2019). A detailed description of the calculation can be found in the original article by Wink et al. (2012). Applying fastECM, we analyzed node centrality on the whole brain level by treating each voxel as a node. Major advantages of EC compared to other rsfMRI analysis methods are its faster computation times, and the fact that it is data driven and does not require additional assumptions (Wink et al, 2012).

Applying the fastECM Algorithm by Wink et al. (2012), we analyzed node centrality on the whole brain level by treating each voxel as a node. For the analyses, voxel-wise EC maps at each time point were compared between groups to infer connectivity changes as a result of sequence-specific motor learning. For example, the changes between d1 and d2 of the LRN group were compared to the changes between d1 and d2 of the SMP group in order to identify potential interaction effects. The comparisons between time points were interpreted as being reflective of the different learning stages. Based on the motor learning literature, we defined fast learning as changes between the first and second days of training (d1/d2), slow learning as improvements between the second and fifth days (d2/d5), overall learning from no knowledge of the task up until plateau (d1/d5) and retention as skill recall after the passage of time without training (d5/d17) (Dayan and Cohen, 2011).

### Behavioral Analysis

We analyzed the behavioral data in order to characterize performance and contextualize our fMRI results within the different stages of sequence learning introduced by task-based fMRI studies. Performance was assessed by measuring temporal synchronization (SYN) and lag-aligned root mean squared error (RMSE). SYN describes the deviation of the FOR bar from the REF bar, in milliseconds (LRN sequence or SMP sequence). A cross-correlation between REF and FOR patterns was computed for each trial, and the time lag with the maximum correlation value was used to determine the degree of temporal synchronization (SYN), (where a score of 0 denotes perfectly synchronized performance). The lag-aligned values were computed by setting the lag values to 0. The resulting deviation values were square rooted and averaged. The square root was then used to calculate the lag-aligned RMSE. RMSE represents the spatial deviation between the position of REF and the position of FOR adjusted by the participant. To ensure that they did not differ behaviorally at the beginning of training, we conducted t-tests and compared the averages of the first block (three trials) of the SMP sequence in both groups for both SYN and RMSE. Behavioural performance metrics were computed using custom-built MATLAB scripts (available at https://github.com/neuralabc/SPFT).

### Statistical Analyses

In order to understand changes in performance as a result of training, we computed the mean SYN per day for each participant and performed a repeated measures ANOVA with factors day (1-17) and group (LRN vs. SMP). Mauchly’s tests were conducted to correct sphericity when necessary (Greenhouse-Geisser if ε < 0.75 or Huynh Feldt if ε > 0.75). Post-hoc Tukey’s tests were used to assess timing-specific significant effects between consecutive days. Statistical testing on the behavioral data was performed with the Jamovi software (https://www.jamovi.org; Singmann, 2018; Lenth, 2018) based on R (https://cran.r-project.org/).

### Group X Time Interaction Analyses

Prior to the interaction analysis and to verify that the groups baseline EC maps did not differ on the days before training, we applied independent t-tests in SPM to compare both groups at d0 as well as d1.

For the rsFC whole brain interaction analyses we used a flexible factorial design for longitudinal data from the CAT12 Toolbox in SPM with two groups and 5 time points (d0, d1, d2, d5, d17). The flexible factorial design included two factors: group and time point. Within the interaction analyses, changes between time points within-group were compared across groups and learning stages. For example, when investigating the fast learning stage, the interaction analysis consisted of comparing both the d1-d2 contrast (decreases from d1 to d2, relative to LRN) and the d2-d1 contrast (increases from d1 to d2, relative to LRN). Interaction analyses were performed for the following time point contrasts: d1-d2, d2-d1, d2-d5, d5-d2, d1-d5, d5-d1, d5-d17, and d17-d5. Following the identification of interaction effects, we then computed the magnitude of change within the significant ROI in each group to determine which group was driving the effect. Based on our design, the identification of a change in centrality that was greater in the LRN group than the SMP group was considered as a sequence-specific effect. RsfMRI results are reported using cluster inference with the SPM default primary threshold of p < 0.001 and FDR correction at the cluster level at p < 0.05 (Woo et al., 2014).

## Results

### Behavioral Results

Two outliers were excluded from the behavioral data because their values were more than 2 standard deviations away from the mean on two days.

There were no differences between the groups on the averages of the first 3 trials of the SMP task in either SYN (t = -1.84, p= 0.07) or RMSE (t = -0.99, p = 0.33).

For the analysis of SYN, we found a significant main effect of group (F(2,57) = 60.2, p < .001, η2 = .68) (Fig. 2). Post-hoc Tukey t-tests revealed significant differences in the LRN group between d1 and d2 (t= 7.89, p <.001) and d2 and d3 (t=4.23, p =.004). There were no significant differences in the LRN group between days 3, 4, 5 and 17. There were no significant differences between days in the SMP group, supporting the hypothesis that the SMP group was not improving in temporal accuracy. Therefore, we were able to see the progress of learning the LRN sequence over time (Fig. 2) by assessing SYN. For the analysis of RMSE, we found a significant main effect of group (F(2,56) = 4.90, p = 0.011, η2 = .149) (Fig.3). Post-hoc Tukey t-tests revealed significant differences in the LRN group between d1 and d2 (t = 4.89, p < 0.001) and d2 and d3 (t = 4.53, p = 0.001). We also found significant differences in the SMP group between d1 and d2 (t = 5.89, p < .001) but no significant differences between d3, d4, d5 and d17 providing evidence for performance improvements during the fast learning phase in the SMP group in spatial accuracy.

**Fig. 2.**
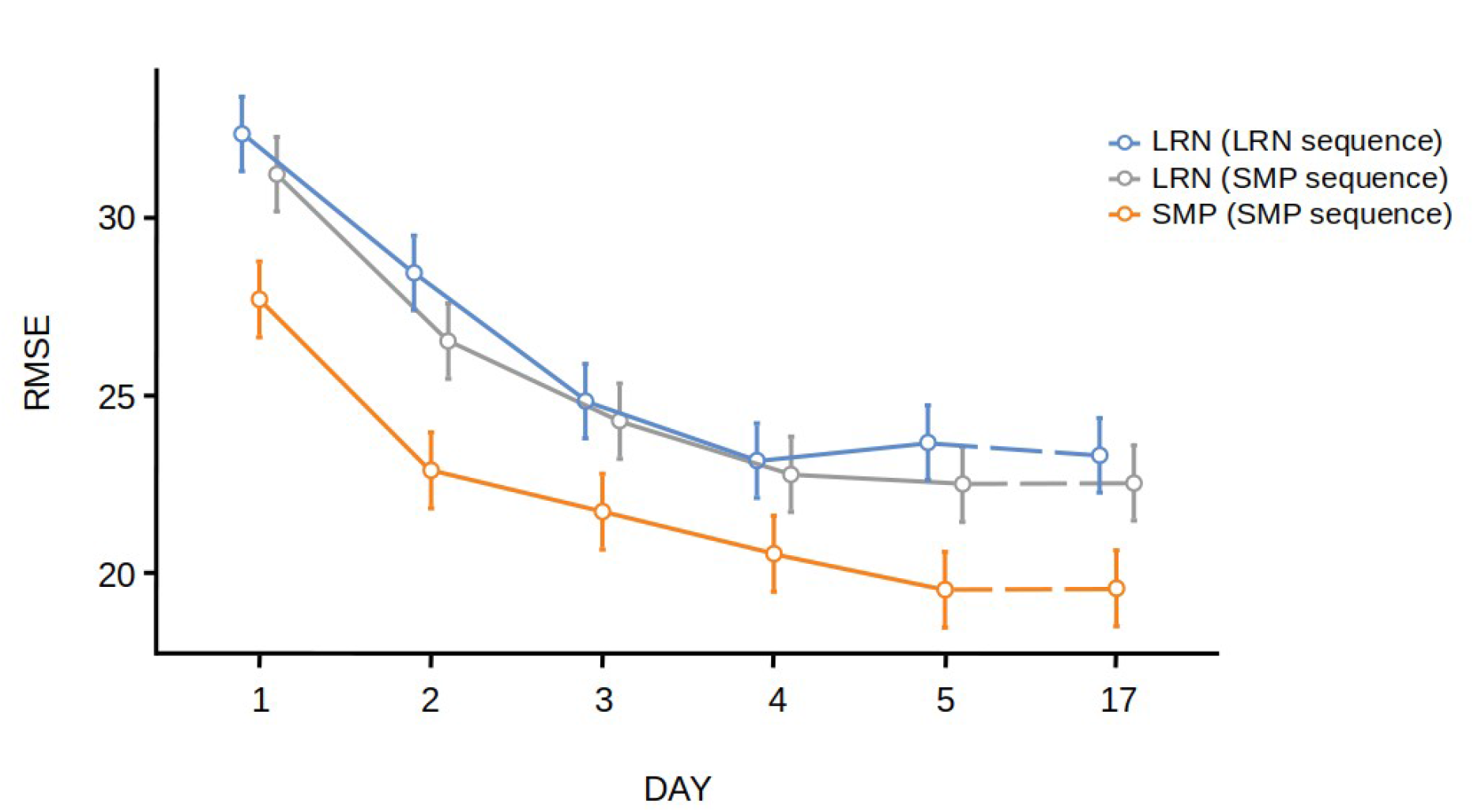
Behavioral Results. Root-mean-squared error (RMSE) for both groups (LRN and SMP) and sequences across all days (d1-d17). Error bars indicate the standard error of the mean. LRN and LRN_SMP values were calculated by averaging 3×3 trials per block. SMP values were calculated by averaging 6×3 trials per block.

**Fig. 3.**
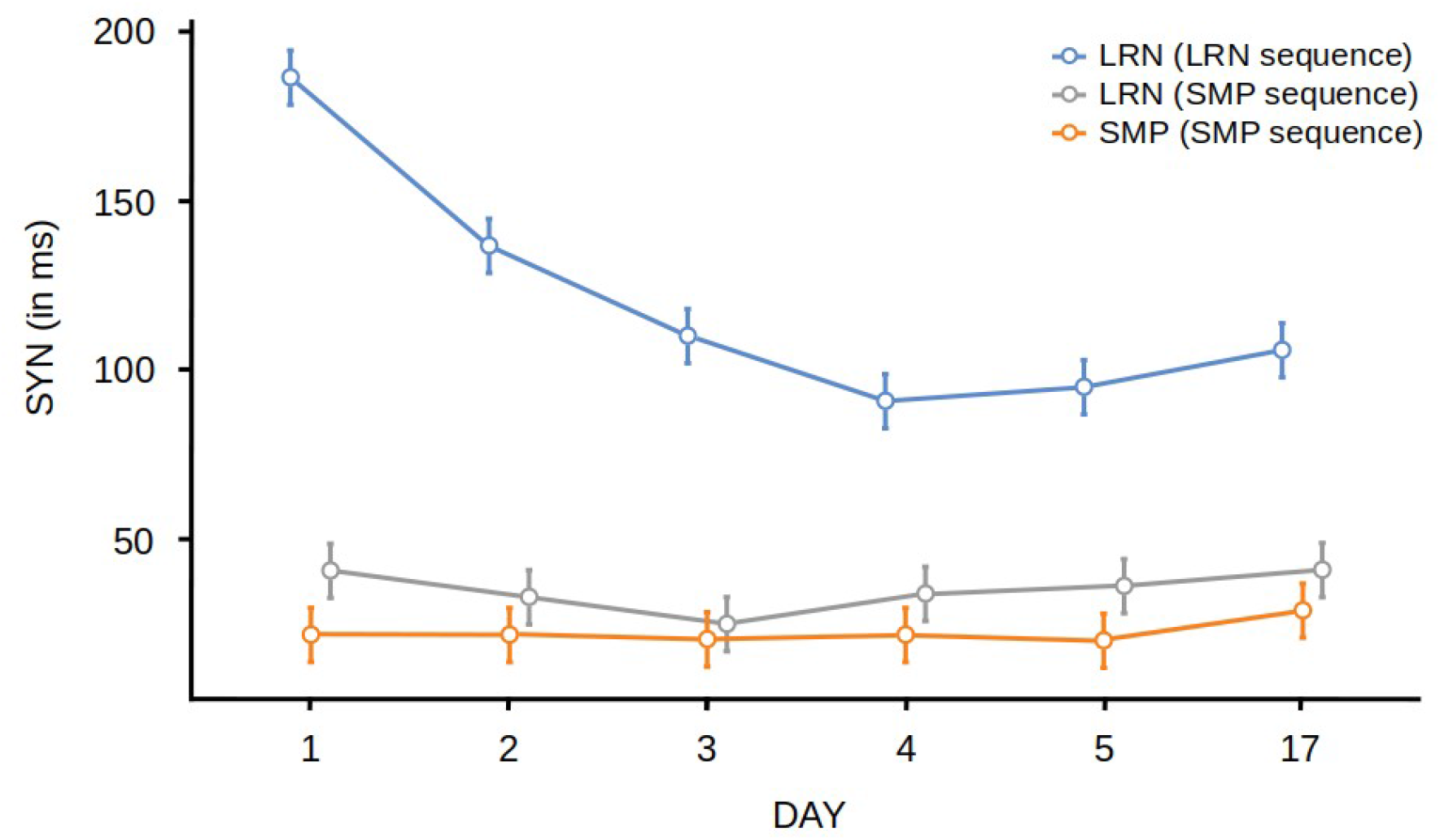
Behavioral Results. Temporal synchronisation (SYN) for for both groups (LRN and SMP) and sequences across all days (d1-d17). Error bars indicate the standard error of the mean. LRN and LRN_SMP values were calculated by averaging 3×3 trials per block. SMP values were calculated by averaging 6×3 trials per block.

There were no significant differences between the two groups in musical and physical exercise experience (all p-values > .05). There were no significant differences in the LRN group between days 3, 4, 5 and 17.

### rsFC Results

There were no significant between-group differences in whole brain EC on either d0 or d1.

### Interaction effects - Fast Learning

When investigating changes in EC we found significant group by day interaction during fast learning in the right anterior insular cortex (AIC) and the right SPC (Fig. 3c & d). Trajectories of change in EC were plotted over all training days in significant clusters of the interaction analyses to provide additional context about the progression of changes within those regions across the time points (Fig. 3b and 3e).

### Interaction effects - Slow Learning

Interaction effects between d2 and d5 were found in EC between groups in the right AIC (Fig. 4a). However, this cluster was not sequence-specific, as the SMP group was driving the changes.

**Figure 4:**
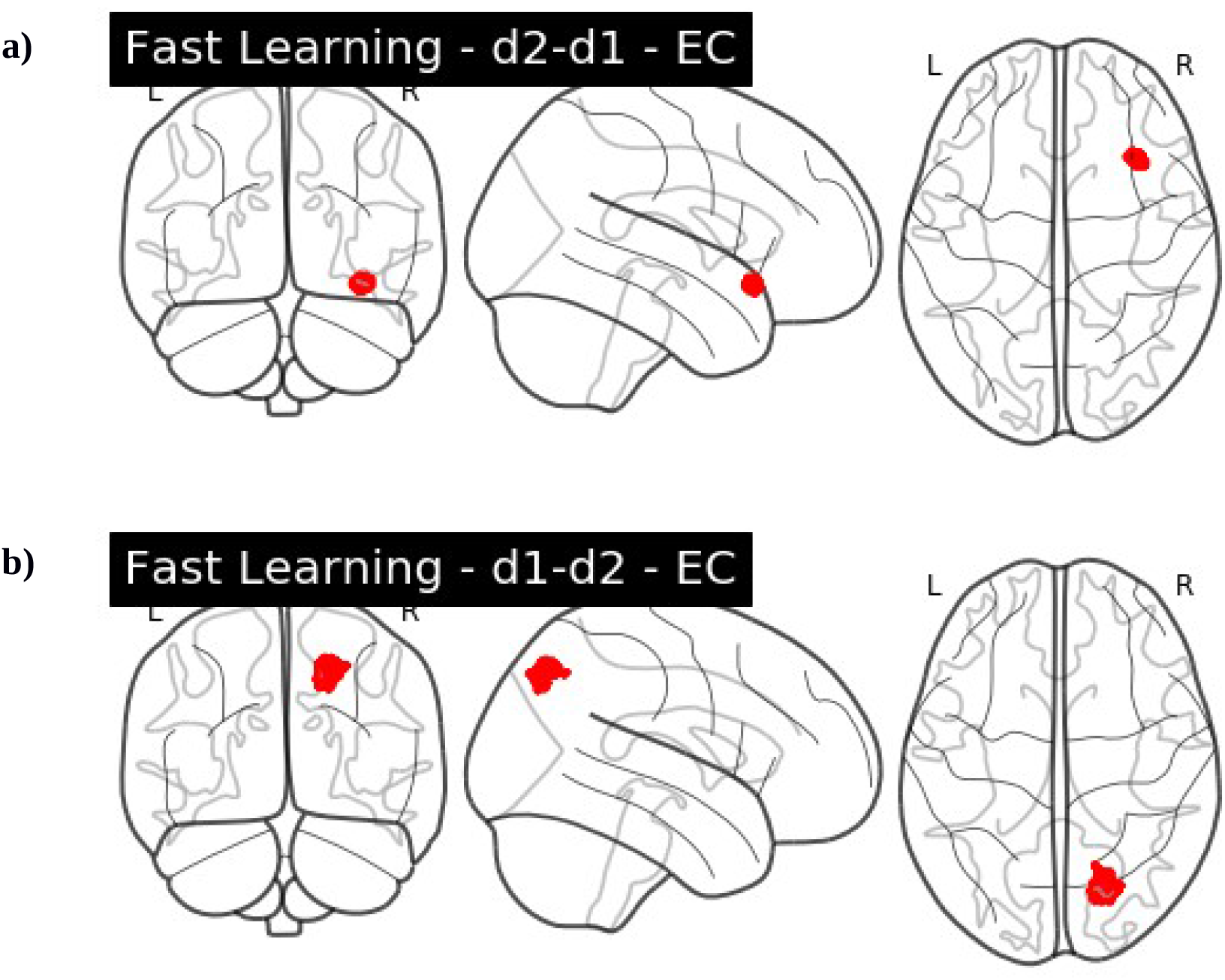
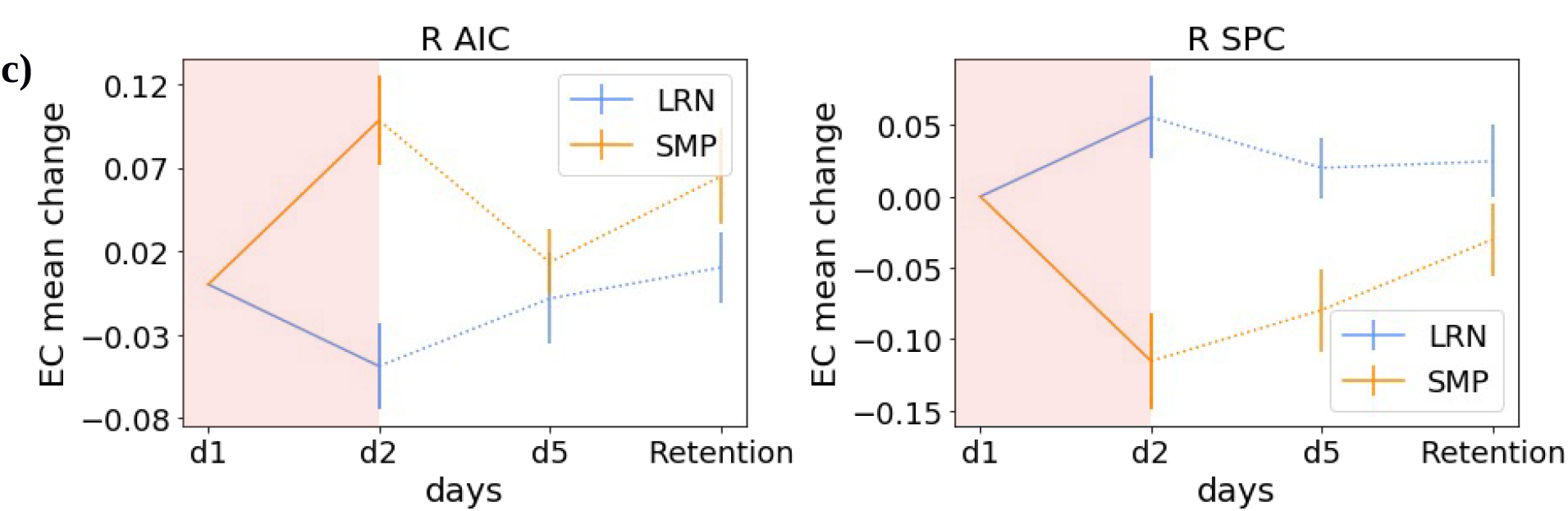
Group interaction: Fast learning. a) Glass brain EC. d1-d2 interaction effect in the right AIC b) Glass brain EC. d2-d1 interaction effect in the right SPC. c) shows the EC change value trajectory over the entire training period for the significant clusters including R AIC and R SPC displayed for LRN and SMP. The period which had the significant interaction (d1/d2) is highlighted in red.

### Interaction effects - Overall Learning

We found interaction effects between groups in EC during overall learning (d1 vs d5) in the right supplementary motor area (SMA) and right parietal operculum (PO) (secondary somatosensory cortex) (Fig. 5c) and bilateral SPC (Fig. 5d). Interaction effects in the right SMA were driven by the LRN group (see Table 4) and therefore identified as sequence-specific.

**Fig. 5.**
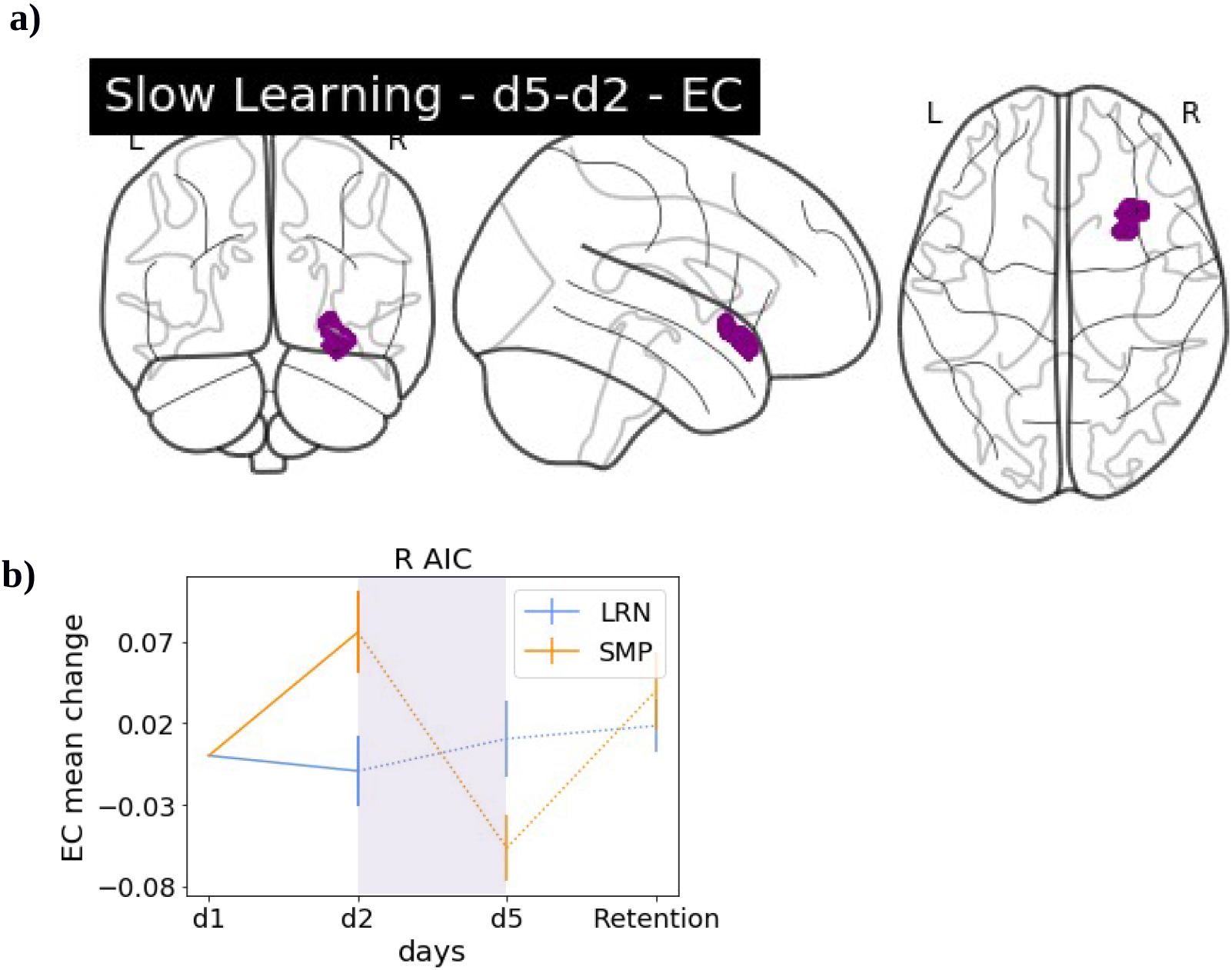
Group interaction: Slow learning. a) Glass brain EC. d5-d2 interaction effects in the R AIC. b) shows the EC change value trajectory over the entire training period for the significant cluster in R AIC displayed for LRN and SMP with the period which had the significant interaction (d2/d5) highlighted in purple

### **I**nteraction effects – Retention

To identify potential degree centrality changes between the last time point of training and the retention probe, we compared d5 and d17 between the two groups. We identified a significant interaction effect in the right Putamen in EC (Fig. 6c). This effect was not predominantly driven by LRN.

**Fig. 6.**
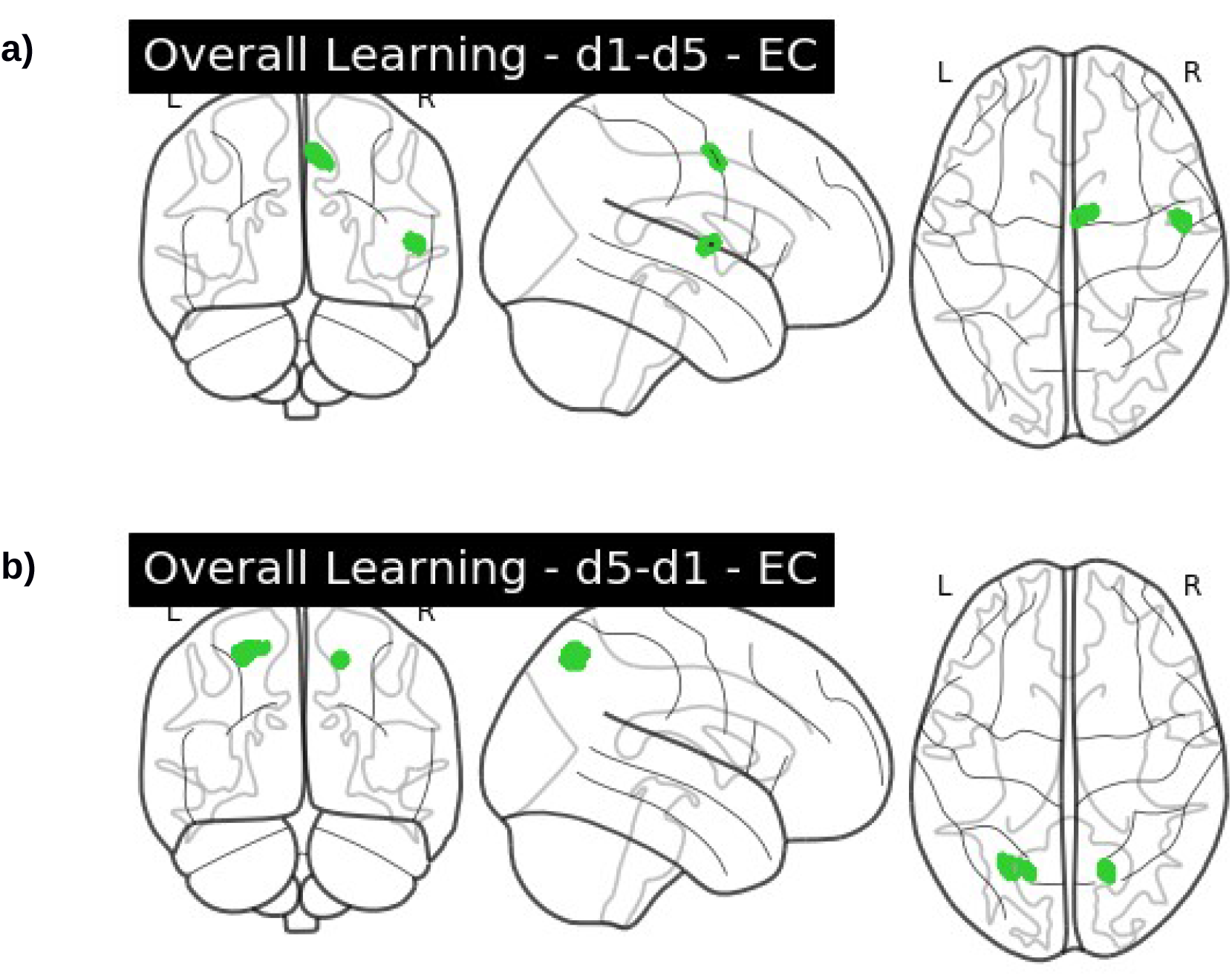
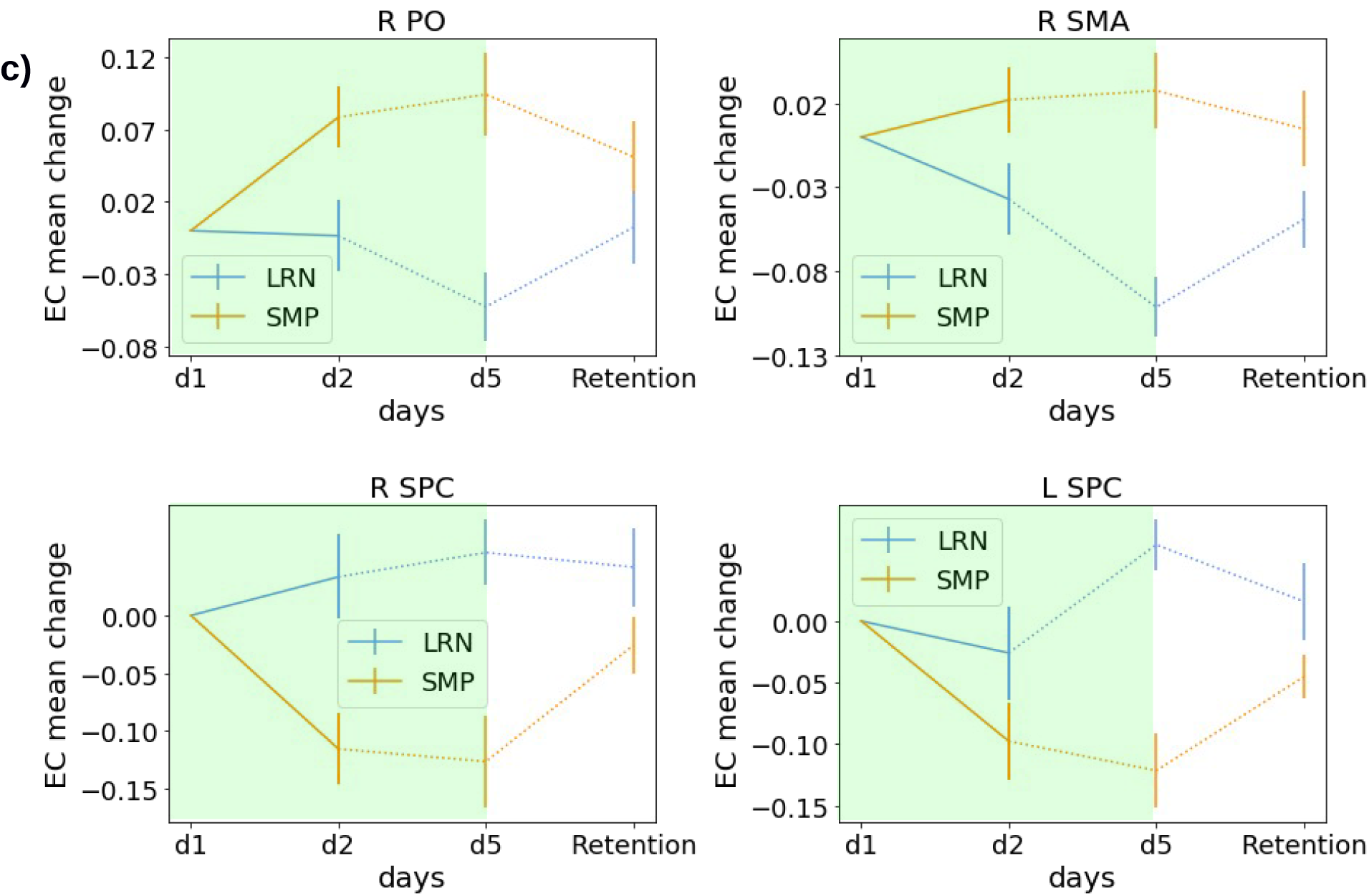
Group interaction: Overall learning. a) Glass brain EC. d1-d5 interaction effects in the R PO and R SMA (seq-spec.). d) shows the EC change value trajectory over the entire training period for the significant cluster in R AIC displayed for LRN and SMP with the period which had the significant interaction (d2/d5) highlighted in green

**Fig. 7.**
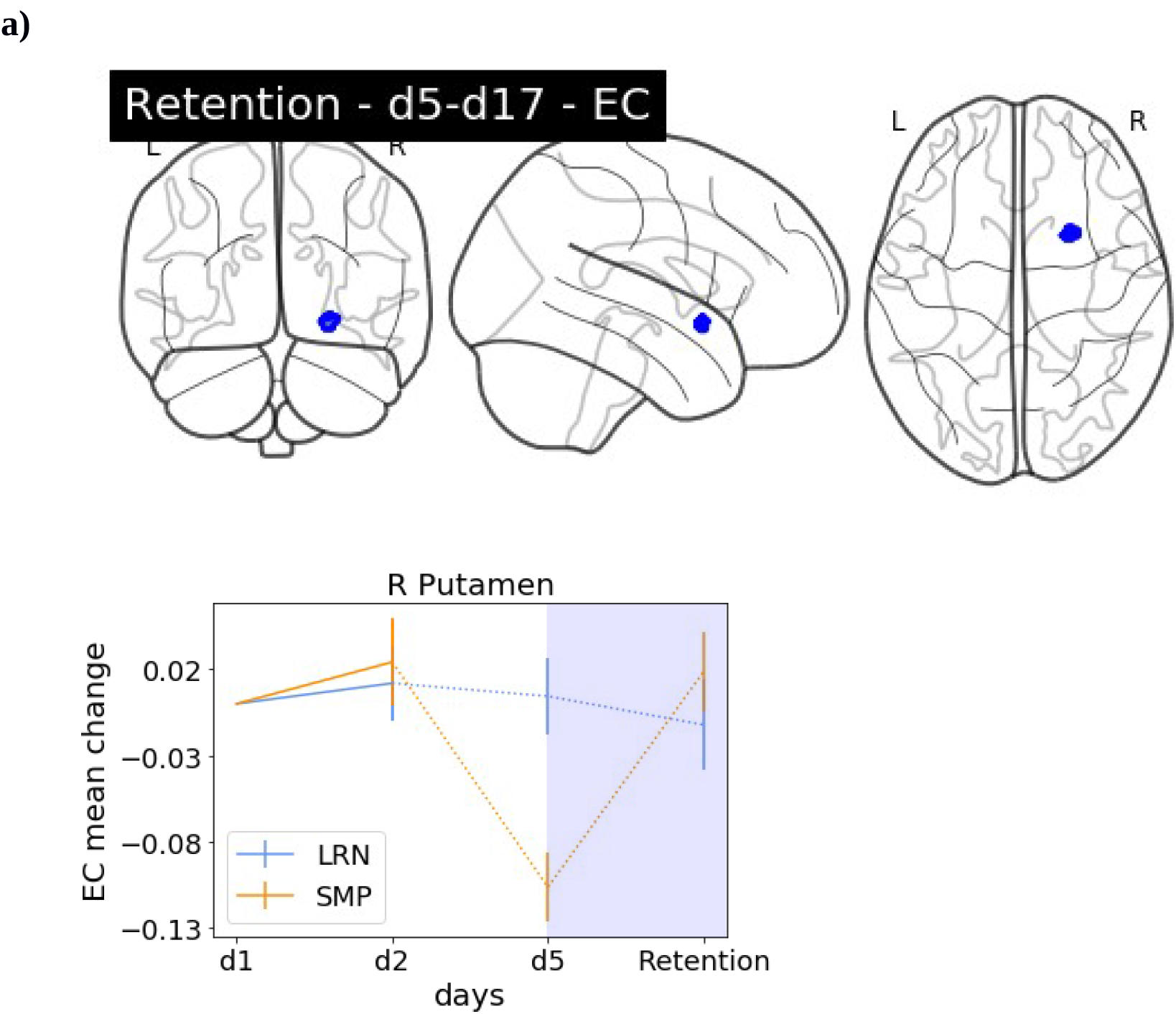
Group interaction: Retention. a) Glass brain EC. d17-d5 interaction effects in the R Putamen d) shows the EC change value trajectory over the entire training period for the significant cluster in R AIC displayed for LRN and SMP with the period which had the significant interaction (d5/d17) highlighted in blue

**Fig. 8.**
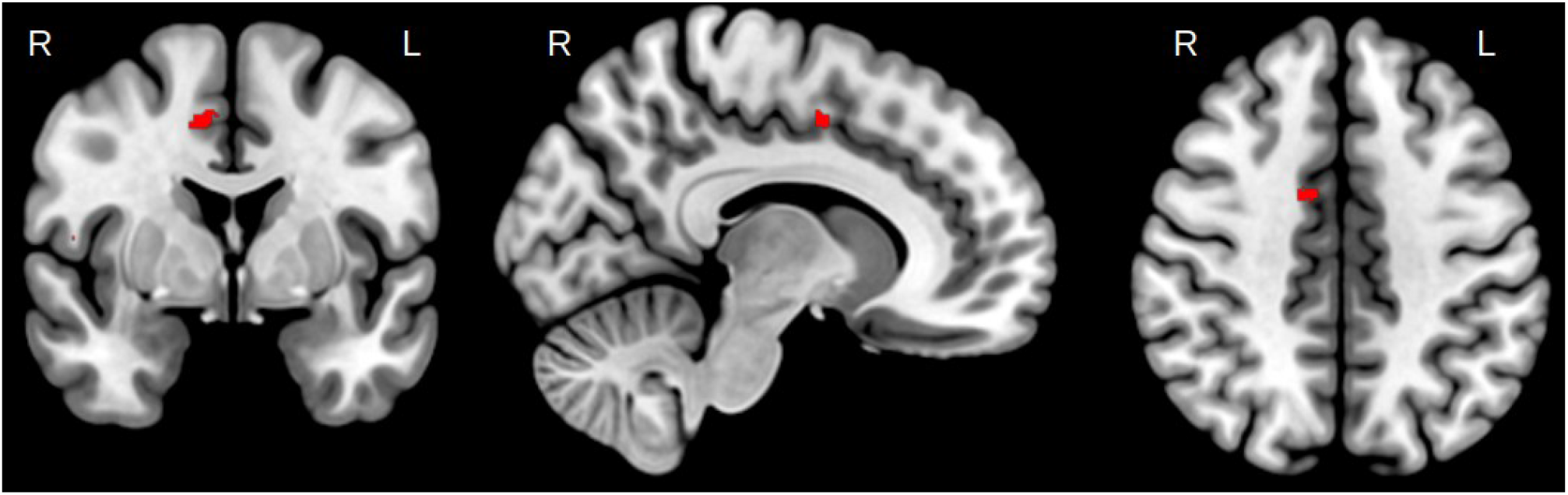
Anatomical location of the sequence-specific cluster in the R SMA

Precise locations as well as MNI coordinates of peak voxels and significance thresholds of the significant clusters are listed in Table 1.

**Table 1:** Flexible Factorial Analysis interaction effect results, EC

| **Contrast** | **pval FDR** | **K** | **peak**<br>**T-value** | **MNI Coordinates** | **Anatomical location** |
| --- | --- | --- | --- | --- | --- |
| d1-d2 | 0.001 | 144 | 4.21 | 35,22, -14 | R AIC |
| d2-d1 | 0.003 | 431 | 3.88 | 24, -76, 47 | R SPC |
| d5-d2 | 0 | 391 | 4.5 | 30, 18, -14 | R AIC |
| d17-d5 | 0.025 | 27 | 4.25 | 24,10,-10 | R Putamen |
| d1-d5 | 0 | 131 | 4.03 | 6,-4,52 | R SMA |
| d1-d5 | 0.002 | 92 | 3.8 | 52,-2,7 | R PO |
| d5-d1 | 0.008 | 244 | 4.47 | -26,-66,55 | L SPC |
| d5-d1 | 0.008 | 88 | 3.81 | 17,-65,52 | R SPC |

**Table 2:**
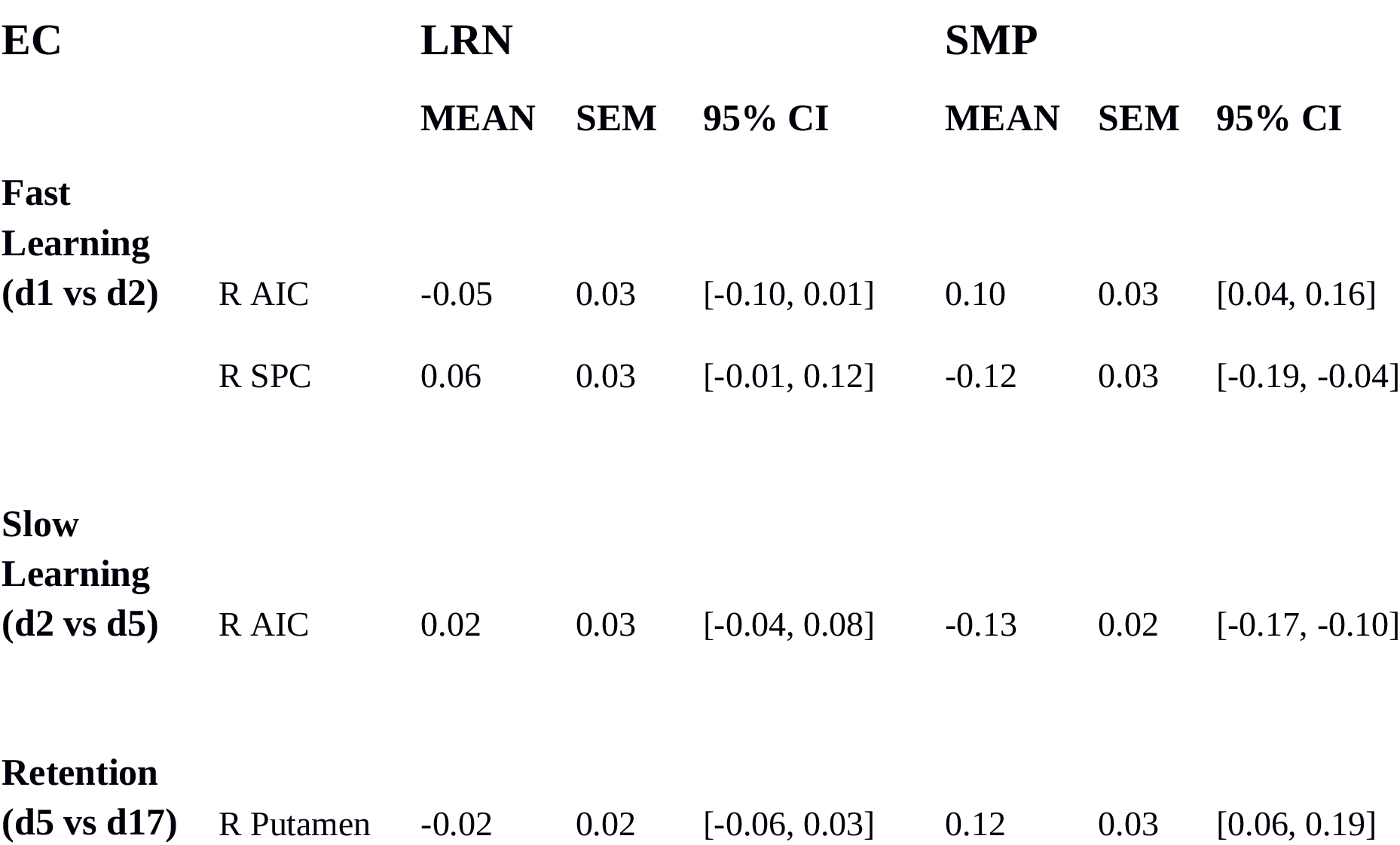
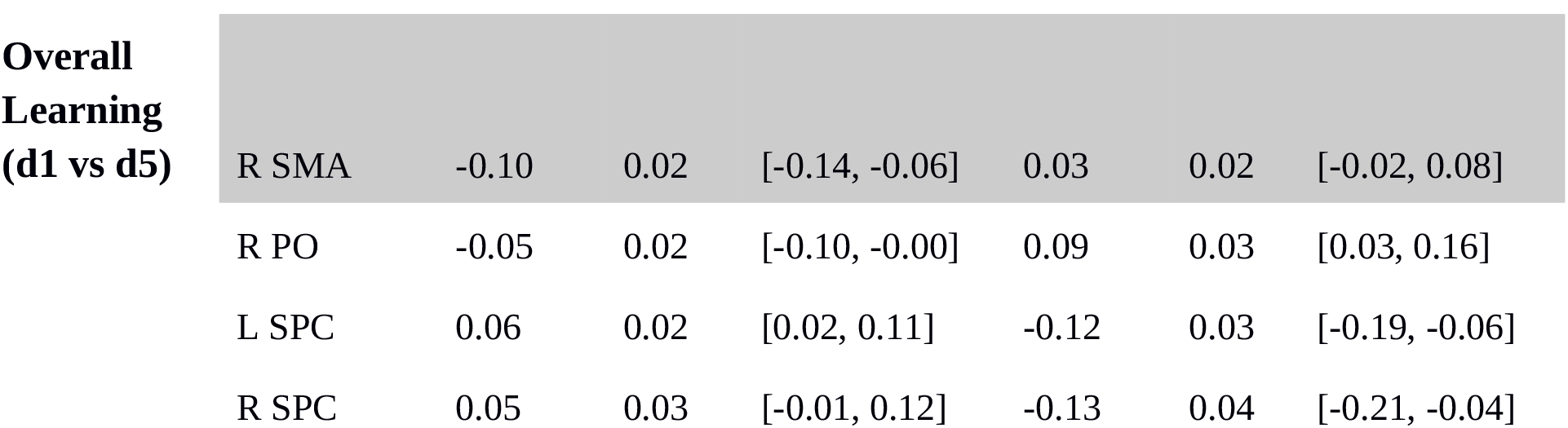

## DISCUSSION

We investigated between-group interaction effects in functional connectivity changes over the course of one week of training on a continuous motor sequence task with rsfMRI. Our study encompasses fast learning, slow learning, overall learning, and retention. Crucially, we compared a group that learned a complex sequence with a control group that performed a matched motor task to distinguish changes due to sequence-specific learning and provide additional context for changes in resting state functional connectivity after MSL throughout the learning stages.

Interaction analyses revealed a set of regions mostly within the motor network that exhibited differential change between the two groups, providing evidence that these regions play a role in functional plasticity after MSL. Our criterion for assigning sequence-specific function to a region was dependent on that region exhibiting greater centrality change in the LRN group. As a result, we found sequence-specific functional changes in one brain region at one time point: the SMA during overall learning. Most of the other regions identified in the interactional analyses, including well known motor learning related areas such as the SPC also showed opposing centrality changes in both groups but were not being driven by LRN, suggesting differential involvement of this region when learning a complex motor sequence versus a simpler motor execution.

### Sequence-specificity

Behaviorally, we were able to show sequence-specific increases in performance over the course of training. We found that the experimental group improved in temporal accuracy over time, while the control group did not, and that the experimental group reached a learning plateau on the fourth day of training. In order to find out which regions were relevant for sequence-specific learning and temporal accuracy, we evaluated which effects in the interaction analyses were 1. driven by the LRN group and 2. showed little to no change in the SMP group. By doing so, we ensured that we were evaluating results in the LRN group reflecting improvements in temporal accuracy.

Hence, functional connectivity changes reflecting sequence-specific learning effects were identified during the overall learning period in the SMA. We found no evidence for sequence-specific effects during the fast-, slow learning or retention phases. A predominant role of the SMA in sequence-specificity during offline learning aligns well with the literature on sequence-specificity from task-based online assessments of MSL (,, Elsinger et al., 2006, Gaymard et al., 1990, Gerloff et al., 1997,, Hikosaka et al., 1999, Jenkins et al., 1994, Lee & Quessy, 2003, Mushiake et al., 1991,, Shima & Tanji, 2000, Tanaka et al., 2010,, Vollmann et al., 2013).

Previous research has shown that the SMA is also relevant for sequence-specific MSL during online learningin nonhuman primates where it has been established that SMA is activated during the planning period prior to sequence execution (Tanji & Shima, 1994).

While it seems counter-intuitive to see such lateralized results on the right side when training on a motor task performed with the right hand, it is important to consider that the interaction analysis shows results only in regions differing between the experimental and control groups. Since both groups are manipulating the device with their right hand, any activity related to simple motor execution will overlap in both groups and, in the case of simple motor movements of the right hand, be present in the left hemishpehre. Therefore, sequence-specific differences between groups may be more difficult to detect in these regions as they may end up being masked by connectivity changes attributable to motor execution. Additionally, previous research has shown that specifically the right SMA is highly connected to other bilateral regions such as the basal ganglia, insula, thalamus and cerebellum, and thus represents a strong network-level affiliation with the entire motor network(Narayana et al., 2012). The right SMA has been found to be relevant for sequential learning (Lacourse et al., 2005, Lin et al., 2012, Mallol et al., 2007, Wymbs & Grafton, 2015), the and specific to the linking of action phases during sequential movements (Säfström & Domellöf, 2018), action execution (Narayana et al., 2012) as well as imagining rhythmic motor execution (Oullier et al., 2005). Our results provide support for the role of the SMA in processing sequential information and illustrate how this region is involved in sequence-specific plasticity offline as a result of MSL. We found that EC in the SMA gradually decreased over the entire learning period, suggesting an ongoing relevance to the learning process. While it would be intuitive to interpret a decrease in connectivity as indicating decreased involvement of that area, it is important to keep in mind that EC is a network measure that reflects less connectedness to other highly connected nodes, rather than less activation of the region per se. A decrease in this case could also point towards a reduction of neural resources as a result of skill-specific efficiency (Wymbs & Grafton, 2015).

### Motor execution

Behaviorally, we were able to show that both groups improved in spatial accuracy. Within our interaction analysis we found significant effects, mostly within the motor network, that were driven by the SMP group in bilateral SPC, possibly reflecting the behavioral improvements in spatial accuracy. The SPC, a key node in the motor learning network (Dayan & Cohen, 2011, Doyon et al., 2018, Krakauer et al., 2019, Penhune & Steele, 2012), showed centrality decreases in SMP. The SPC is involved in learning the association of visual stimuli and motor actions / visuomotor control (Hardwick et al., 2012, Mutha et al., 2011, Müller et al., 2002), and has been found to be relevant for early and later learning stages (Ma et al., 2011). We observed a steady decrease in SMP in functional connectivity in both regions from d1 to d2 and d1 through d5. Considering that the d1/d2 comparison reflects fast learning and the d1/d5 reflects the overall learning process, we propose that the SPC exhibits a slow change in connectivity over the course of the entire learning period resulting in a centrality decrease for SMP. These centrality changes could reflect a continuous learning process that may serve to integrate the presented visual sequence with the required motor response. While it is reasonable to conclude that the SPC is relevant for both LRN and SMP, as the tasks performed by both groups require the coordination of visual and motor information, it is interesting to note that while there are functional changes evident in the LRN group that may be related to the behavioral improvements, they are not as distinct as those in the SMP group.

Interestingly, within our trajectory visualizations, we found that the changes in centrality in most regions showed opposite directionalities. It is possible that these different directionalities of change as well as the differences in magnitude of change could be due to task complexity (Carey et al., 2005, Witt et al., 2008). We speculate that SMP, being a rather simple task, would cause centrality changes to occur faster, such that we are presented with change reflecting a well-learned or overlearned task (motor execution). In contrast, centrality changes in LRN would rather reflect changes during the actual learning process of the complex sequence within our experimental group.

## Conclusion

The present experiment employed a mixed longitudinal training design with two groups (learning and control) to investigate learning on a continuous motor sequence task. We provide evidence for connectivity changes in the right SMA being specific to the learned sequence rather than motor execution.We also argue that changes in resting state centrality in other regions of the motor network, including the SPC, could be due to motor execution processes common to both tasks.

## Declarations

### Funding

This work was supported by the Max Planck Society, the MaxNetAging Research School (Max Planck Institute for Demographic Research, ATJ), the NWO Vici grant (PI: Birte Forstmann)(PLB), the Heart and Stroke Foundation of Canada (CJS, CJG), the Michal and Renata Hornstein Chair in Cardiovascular Imaging (CJG), the National Science and Engineering Research Council (CJS: RGPIN-2020-06812; CJG: RGPIN 2015-04665) and the Fonds Recherche Québécois Nature et Technologies (CJS).

### Conflicts of interest/Competing interests

Not applicable

### Availability of data and material

Statistical maps will be made available upon reasonable request and released on Neurovault following publication. Due to the ethical standards of the Ethics Committee of the University of Leipzig raw data cannot be made available.

### Code availability

The preprocessing pipeline is available at: https://github.com/AthSchmidt/MMPI/tree/master/preprocessing. Behavioural performance metric computation scripts are available at https://github.com/neuralabc/SPFT.

### Ethics approval

All study procedures were in accordance with the general ethical standards of the Ethics Committee of the University of Leipzig and in line with the 1964 Helsinki declaration. The study was further approved by the Ethics Committee of the University of Leipzig.

## Notes

https://github.com/AthSchmidt/MMPI/tree/master/preprocessin

https://github.com/neuralabc/SPFT

### Competing Interest Statement

The authors have declared no competing interest.

### Summary of Updates

We had to omit the previously included DC measures due to a conceptual mistake in the pipeline.

